# Decoding auditory working memory content from EEG aftereffects of auditory-cortical TMS

**DOI:** 10.1101/2024.03.04.583379

**Authors:** Işıl Uluç, Mohammad Daneshzand, Mainak Jas, Parker Kotlarz, Kaisu Lankinen, Jennifer L. Fiedler, Fahimeh Mamashli, Netri Pajankar, Tori Turpin, Lucia Navarro de Lara, Padmavathi Sundaram, Tommi Raij, Aapo Nummenmaa, Jyrki Ahveninen

## Abstract

Working memory (WM), short term maintenance of information for goal directed behavior, is essential to human cognition. Identifying the neural mechanisms supporting WM is a focal point of neuroscientific research. One prominent theory hypothesizes that WM content is carried in “activity-silent” brain states involving short-term synaptic changes. Information carried in such brain states could be decodable from content-specific changes in responses to unrelated “impulse stimuli”. Here, we used single-pulse transcranial magnetic stimulation (spTMS) as the impulse stimulus and then decoded content maintained in WM from EEG using multivariate pattern analysis (MVPA) with robust non-parametric permutation testing. The decoding accuracy of WM content significantly enhanced after spTMS was delivered to the posterior superior temporal cortex during WM maintenance. Our results show that WM maintenance involves brain states, which are activity silent relative to other intrinsic processes visible in the EEG signal.

## Introduction

Working memory (WM), the brain system that enables maintenance and processing of recent information, plays an essential role in daily living. The mechanisms and brain areas underlying WM maintenance have thus been prominent topics for neuroscience research. However, research into its neuronal mechanisms has resulted in seemingly contradictory results that have led to a long-standing controversy. The prevailing hypothesis suggests that information is maintained through persistent firing in the prefrontal cortex (PFC). Conversely, an alternative theory posits that persistent activity is not necessary for WM maintenance and rather maintenance can be dynamical in an ‘activity-silent’ format via functional connectivity and/or synaptic weights. [1–3]. Much of this research has been conducted in the visual modality only, leaving some of the most ecologically relevant aspects of WM in other sensory modalities relatively underexplored. One such aspect is auditory WM, which enables temporary storage and manipulation of sounds and verbal information, such as spoken language or music.

Initially, WM maintenance was linked to persistent activity of prefrontal neurons that respond to the incoming stimulus and remain activated even after the stimuli have vanished [4–6]. However, subsequent human neuroimaging studies suggested that the content of visual WM could only be decoded from signal-change patterns in sensory and posterior brain areas where persistent activity is not present during WM maintenance [7–9] (however, see also [10–13]). Studies in non-human primates (NHP) have not shown persistent WM related neuronal activity during the maintenance period in sensory areas [14, 15]. The identification of content-specific persistent firing patterns at the sensory level has proven challenging in NHP studies of auditory WM as well [16–18]. At the same time, human studies have managed to decode auditory WM content from fMRI signals [12, 19–21] as well as intracranial EEG signals [22] from auditory cortices. The diverse and contrasting findings have inspired the development of a family of alternative theories suggesting that WM is maintained in a more distributed and dynamic fashion than initially believed [23–25].

In explaining possible mechanisms for ‘activity-silent’ WM maintenance, a theory proposes that WM is maintained through intermittent bursts of neuronal firing and intervals of short-term synaptic plasticity (STSP) [26], i.e., transient changes in the strength of synaptic connections between neurons [27]. Item-specific activation of neurons during the encoding process leads to presynaptic accumulation of calcium, which facilitates postsynaptic connectivity. Due to this calcium buffer, which operates on a time scale of seconds, even sparse bursts of firing will be sufficient to maintain the “activity-silent” WM representations [3, 25, 26]. The network maintaining a synaptic WM trace will respond in a content-specific fashion, even if the non-specific input is completely unrelated to the maintained representation [3]. Hence, information maintained in WM via the activity-silent synaptic mechanisms should be decodable with machine learning techniques that can classify responses elicited to any unrelated stimulus that broadly activates the same neuronal population [3]. To test this prediction, recent human EEG and MEG studies presented participants with “impulse stimuli”, such as strong visual or auditory feature patterns unrelated to the maintained content, during WM maintenance [13, 28–31]. The content maintained in WM, which is otherwise in an activity-silent (or “hidden”) state, became more readily decodable from EEG or MEG responses to such impulse stimuli [13, 28–31]. A limitation in many of these studies, however, is that it is difficult to deliver such impulse stimuli to a particular brain area only.

A non-invasive way to probe hidden brain states that underlie human cognition is transcranial magnetic stimulation (TMS). Unlike observational methods such as MEG or EEG, TMS allows us to causally interact with focal areas whose role in WM we intend to evaluate [32]. In studies of human memory processes, TMS has been used to modulate maintenance of visual WM representations [33] and to enhance neuronal plasticity in visual cortex [34]. A particular benefit of using single-pulse TMS as opposed to task-irrelevant sensory stimuli for probing memory-related brain states that its effects are both temporally and anatomically specific [35]. In a recent study that used TMS to enhance WM decoding from EEG signals [28], perturbing the WM circuits during the seemingly activity-silent maintenance period yielded noteworthy results. This intervention not only augmented the decoding of representations that were stored passively in memory compared to actively maintained content but also contributed to participants recalling passively maintained items more effectively from WM. However, to our knowledge, this approach has so far not been tested in WM studies targeting auditory or other earlier sensory cortex areas, or in designs applying active and sham TMS in the same participants.

Thanks to recent advances in MRI-guided TMS navigation systems and more focal stimulation coils, TMS pulses can be delivered to the area of interest at an exact latency. This allows one to test anatomically and temporally specific hypotheses to develop an understanding of how sensory areas might be contributing to WM. Here, we investigated whether the content of auditory WM, which is embedded in an activity silent brain state, can be decoded from cortical effects of single-pulse TMS, delivered to posterior non-primary auditory cortex during the WM maintenance period. This non-primary auditory cortex target was in in the left posterior superior temporal cortex (pSTC). In our multivariate pattern analysis derived from whole-scalp EEG, the decoding accuracy increases above chance level directly after the TMS pulse in an Active TMS condition, but not after a Control TMS pulse that was too weak to activate the pSTC target area. Therefore, our study provides strong evidence for the activity silent theory of WM maintenance.

## Methods

### Participants

A total of 23 healthy right-handed [36] participants (12 women, 11 men; mean age ± standard deviation, SD, = 32 ± 12 years) were enrolled. One participant was excluded due to excessive movement artifacts (facial movement artifacts in more than 50% of trials) and another due to their chance-level behavioral performance, resulting in a final cohort of 21 participants (11 women, 10 men; mean age ± SD=31±10 years) for the Active TMS session. Significantly better than chance level performance is essential in WM experiments to ensure that task performance is not a result from a mere guessing. As for the Control TMS session, two participants opted not to continue the study and one dataset was rejected due to excessive noise, resulting in a final sample of 18 participants (9 women, 9 men; mean age ± SD = 31 ± 11 years). The same participants attended Active TMS and Control TMS conditions to eliminate variability in EEG responses from different participants in different conditions. All participants had normal or corrected to normal vision and self-reported normal hearing. The participants provided written informed consent and were informed that they could withdraw at any time. A monetary compensation was given for each visit. The study design, protocol, and consent form were approved by the Mass General Brigham Institutional Review Board.

### Stimuli and experimental paradigm

#### Auditory stimuli

We employed non-conceptual, parametrically varied ripple sounds as WM items (**Fig. 1a**). Such stimuli do not allow verbal memorization strategies. The ripple-velocity pool was individualized based on each participant’s pre-determined ripple-velocity discrimination thresholds. Just noticeable difference (JND) values were calculated individually for each participant using a 2-down, 1-up staircase algorithm [12, 13, 22]. Based on these values, we created four auditory ripple sound stimuli, ensuring that each stimulus was positioned 1.5 JNDs apart from its closest neighbor in velocity. We used four different ripple sounds as to-be-remembered stimuli. The same four ripple stimuli were also presented as test stimuli. The order of the memory items was pseudo-randomized. The participants were naïve to the number of memory items. The auditory stimuli were presented at a comfortable listening level through Sensimetrics S14 Insert headphones (Sensimetrics, Malden, MA) that provide high-quality acoustic stimulus delivery while attenuating TMS click noise, analogous to our previous studies [37].

**Figure 1.**
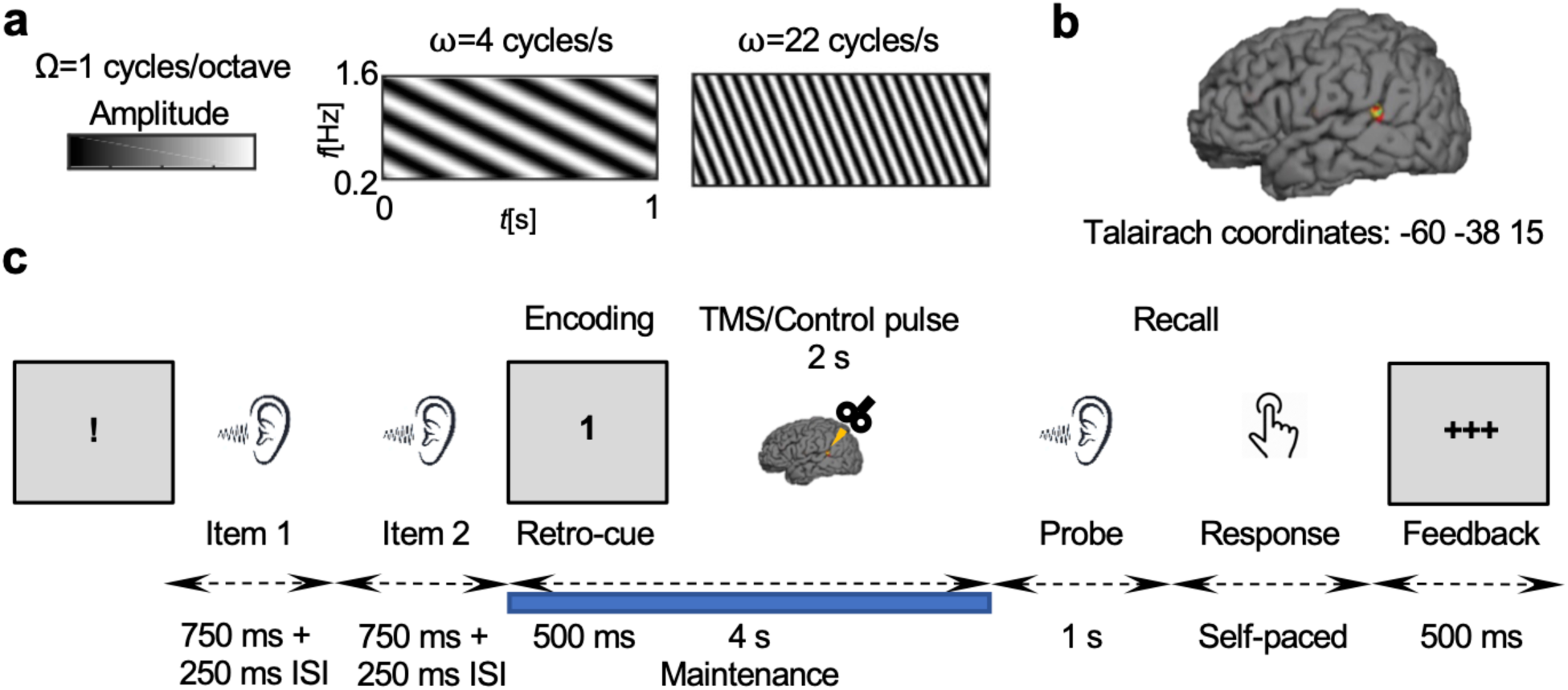
Task design. **(a)** Examples of the modulation patterns for ripple sounds **(b)** The target brain area for the TMS pulse, adapted from Uluç and colleagues [21]. **(c** The auditory WM retro-cue paradigm. The timeline of events in one trial is depicted.

#### Experimental paradigm

Figure 1 shows the retro-cue experimental paradigm used during the TMS-EEG recordings. Each trial started with a “!” presented on the screen. It was followed by two different consecutive ripple sounds (memory items) that were presented for 750 ms with a 250 ms interstimulus interval. Ripple sounds were followed by a visual retro-cue indicating whether the participant had to remember the first sound (”1” on the screen) or the second sound (“2” on the screen). This was followed by a 4-second maintenance period, where a TMS pulse was delivered in the middle. We delivered the TMS pulse at the 2 s mark to have the same amount of signal before and after the TMS pulse during maintenance for balanced comparison of results. Next, a test sound was presented. The task was to determine whether the test sound was the same or different from the memorized item. Participants responded with a mouse click with their right hand. An index finger click indicated that the sounds were the same and a middle finger press indicated that they were different. Finally, the screen showed whether the participant had responded correctly or incorrectly. One run of the experiment consisted of 48 trials and one session consisted of 6 runs. Thus, one session had 288 runs in total.

### Structural MRI Data Acquisition

T1-weighted anatomical images were acquired for with a multi-echo MPRAGE pulse sequence (TR=2510 ms; 4 echoes with TEs=1.64 ms, 3.5 ms, 5.36 ms, and 7.22 ms; 176 or 208 (to cover the ears) sagittal slices with 1×1×1 mm^3^ voxels, 256×256 mm^2^ matrix; flip angle = 7°) [38] in a 3T Siemens Trio MRI scanner (Siemens Medical Systems, Erlangen, Germany) using a 32-channel head coil.

### TMS-EEG Data Acquisition

To be used as stimulus amplitude, resting motor threshold (rMT) of each participant was measured by sending a pulse to the left motor cortex thumb area and measuring the response from first dorsal interosseous muscle of the dominant right hand. From peak to peak, the smallest stimulation intensity resulting in 5/10 responses with amplitudes was equal to or greater than 50 uV. After the the rMT visit, participants completed two single-blind sessions. TMS pulses were delivered either 1) at 100% of individual rMT to the posterior nonprimary auditory area pSTC in the left hemisphere (“Active TMS”) at 45° angle relative to the reference vector [0 0 −1] (A 0-degree rotation relative to this reference vector means the coil handle is oriented from anterior to posterior. The rotation angle increases counterclockwise around the superior-inferior axis) or 2) at 100% of rMT at the same location and same angle but with a 20 mm plastic block between the coil and scalp (“Control TMS”). The pulses were delivered 2 s into the 4 s maintenance period. EEG, horizontal EOG, and ECG data were sampled at 25 kHz with a 64-channel active EEG system (ActiChamp, Brain Products GmbH, Gilching, Germany). TMS was delivered with a MagPro X100 w/ MagOption stimulator and a C-B60 figure-of-eight coil (MagVenture, Farum, Denmark). The plastic block used in the control sessions was built in-house and was the same shape as the TMS coil. The order of Active and Control TMS sessions was counterbalanced across the participants.

In both Active TMS and Control TMS conditions, the TMS coil clicks were masked with 8 kHz low pass filtered white noise throughout the experiment. The white noise and the sound stimuli were presented through Sensimetrics S14 Insert headphones (Sensimetrics, Malden, MA) with Comply Canal In-Ear Tips (Hearing Components, Inc., North Oakdale, MN) that have a Noise Reduction Rating (NRR) of above 29 dB. The sound level of the noise mask was measured using Larson Davis sound level meter LXT2 with a Larson Davis RA0038 coupler (Larson Davis, New York, NY): The level of the white noise was at 73 dB SPL and auditory items was at 86 dB SPL. Additionally, subjective report from each participant was taken that they did not hear the TMS clicks or other background noise.

### TMS Neuronavigation

Continuous recording of the head position and orientation relative to the TMS coil was achieved through a commercial TMS neuronavigation system (LOCALITE GmbH, Bonn, Germany) with an optical camera and passive trackers (Polaris Spectra, Northern Digital Inc., Waterloo, Ontario). The participant’s registration to their anatomical data were all done in the Localite neuronavigation software. The reconstructed MRI images were used in the Localite neuronavigation system (LOCALITE GmbH, Germany) to guide the TMS procedure with MRI.

### E-field Calculation

Data from one participant were discarded due to technical problems for Active TMS and Control TMS sessions. To confirm that we had stimulated the intended cortical target, we computed the TMS-induced Electric fields (E-field) using the Boundary Element Method accelerated by Fast Multipole method (BEM-FMM) MATLAB toolbox implementation [39, 40]. The TMS coil locations/orientations were extracted from the TMS navigation software. The participant-specific anatomically realistic high-resolution head models were generated from the T1-weighted images using the SimNIBS toolbox [41]. The model included five distinct layers of scalp, skull, cerebro-spinal fluid (CSF), grey matter, and white matter, with the assumption of uniform conductivity within each layer. The E-fields were calculated on a cortical surface halfway between the grey and white matter surfaces. For group-level visualization, the individual E-field maps were resampled to the FreeSurfer template brain fsaverage (version 6.2) and averaged across participants [42].

### Basic EEG Preprocessing and Analysis

EEG was preprocessed using MNE Python [43]. We used an established, rigorous preprocessing procedure [44, 45]. The data were first detrended, and after selection and interpolation of noisy channels (on average 4, channels), they were epoched to exclude any potential TMS pulse artifacts. Two consecutive ICAs were calculated for the concatenated epochs [44, 45]. The first ICA was used to remove remaining TMS related artifacts (3 independent components were removed), and then the second ICA was performed to exclude physiological artifacts (on average, 5 independent components). Afterwards, we applied a 60-Hz notch filter to remove line noise and a 150 Hz low pass filter. The data were then downsampled to 1 kHz. We did not use any high pass filtering as it might introduce artifacts in the signal and tamper with later TMS-evoked potential components [46]. Finally, all epochs were visually inspected for remaining artifacts and noisy epochs were rejected. Signals from the occipital Iz electrode were excluded from all analyses due to excessive noise in most of the datasets.

To display the time course of brain activity, we calculated event-related potentials (ERPs) separately for each memory item during the memory period as well as a grand average for the whole trial. For memory item comparison, the cleaned data were separated into four groups according to cued memory item. Then, the separated data from all runs were concatenated and averaged across all trials for each memory item within each participant. The data were then averaged across all participants per memory item. For grand averages, the averages were calculated across all trials irrespective of memory item. Topographical maps were calculated with a time window of 0.5 s with equal weights for all trials.

### Cross-participant Multivariate Pattern Analysis

**Figure 2** shows the MVPA pipeline for the cross-participant classification analysis. We conducted the decoding analysis by employing the support vector machine (SVM) implementation from libsvm [47] as provided in the MATLAB/Octave CoSMoMVPA package [48]. To help generalize the analysis results to a larger population, we performed the classification across the participants [13]. For the MVPA analysis, to ensure that any TMS artifacts did not bias the results, the time window starting 50 ms before and ending 50 ms after the TMS pulse was excluded from the analysis. Using CoSMoMVPA and Fieldtrip toolboxes in each participant, we first balanced the number of trials for each class and then calculated the class-specific averages for each participant. Next, for spatial feature selection, we used principal component analysis (PCA) in MATLAB to transform the data to “virtual channels”, extracting the first eight principal components (PC) (**Fig. 2**). To keep the dimensionality constant to allow cross-validation, we selected the number of PCs based on the grand average of cumulative variance explained across all participants, conditions, and WM classes. The number of selected PCs refers to the point where the slope of the tangent decreased below one in a normalized plot with both dimensions scaled between zero and one.

**Figure 2.**
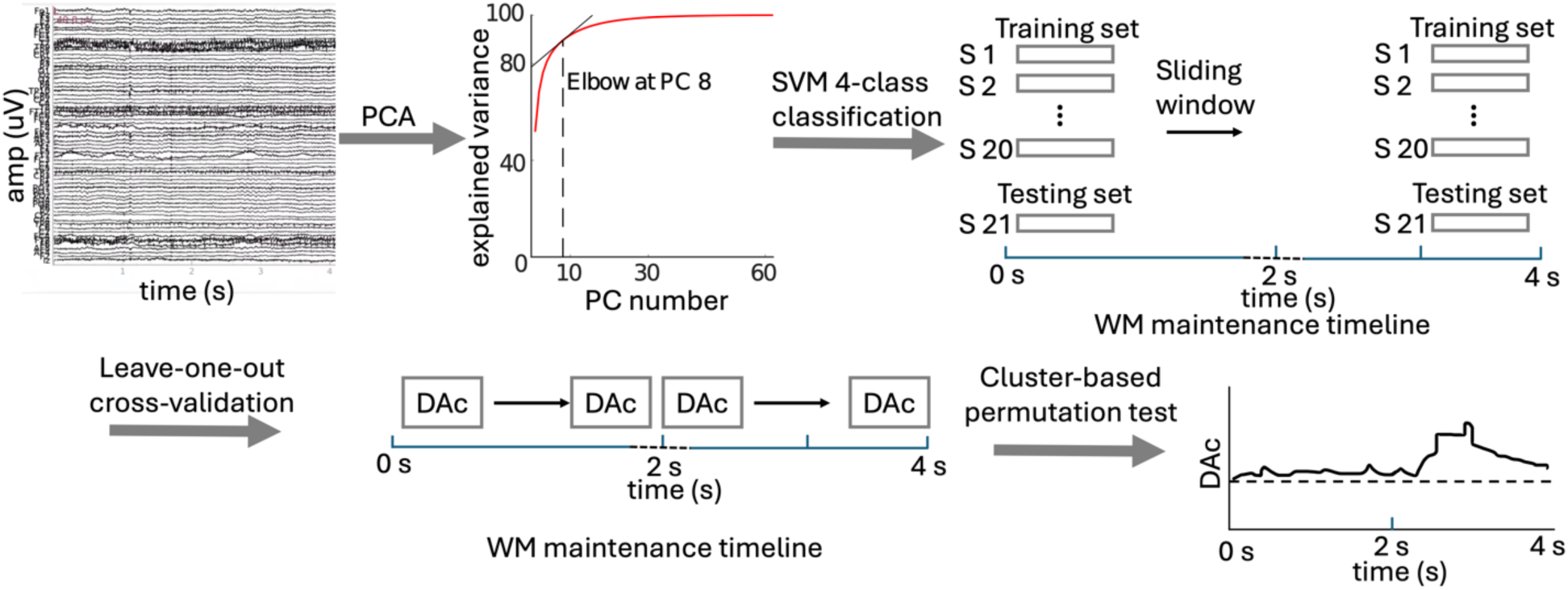
MVPA pipeline. Preprocessed whole head EEG data was entered into a PCA for spatial feature selection. The cut-off for the PC selection (*n*_PC_=8) was determined based on the elbow in the grand-average cumulative variance curve, calculated across all conditions, WM classes, and participants. Then the data was entered into a searchlight analysis with cross-participant 4-class SVM classification. Leave-one-out method was used for cross-validation. The analysis resulted with a decoding accuracy (DAc) time series where DAc is the value assigned to the centroid of searchlight sliding window. For statistical significance, we used maximum statistics with 500 permutations.

Calculating these spatial PCAs separately for each class in each participant ensured that no leakage of information occurred between participants/classes. In each task condition, including periods before and after the pulse in the TMS and Control conditions, this resulted in a two-dimensional (*N*_*subjects*_ × 4) × 8 feature matrix that was entered into the SVM (in the TMS conditions *N*_*subjects*_ = 21, in the control conditions *N*_*subjects*_ = 18).

The classification was conducted as a temporal searchlight analysis [48] with a 300 ms sliding window at 3 ms steps, and was done separately for the periods before and after the TMS pulse. The classification was performed using a leave-one out cross-validation procedure: the data sets were partitioned to training and test sets such that the class-specific sub-averages of one participant used as the test set and those from the rest of the participants as the training set. The decoding accuracy was averaged across all iterations. For each condition (active TMS, Control TMS) and maintenance period (before or after the TMS pulse), the analyses resulted in time series with decoding accuracies of each searchlight centroid (**Fig. 2**).

### Statistical Significance, Cross-participant MVPA

In our cross-participant MVPA approach, the data of one participant were, iteratively, used as the test set, to evaluate the model trained in the other participants. This analysis yields one decoding accuracy value for the entire group at each time point. Instead of conventional one-sample t-tests, we therefore determined the statistical significance of our cross-participant decoding results using robust cluster-based permutation testing, which handles multiple comparison problems using a *maximum-statistic* strategy [13]. Analogous temporal cluster-based maximum-statistic approaches have been widely used procedures in univariate analyses of ERP and MEG data [49]. In this procedure, we first generated 500 unique permutations of the true item-content labels of the classifier. The temporal searchlight analysis was repeated with these permuted labels to generate a distribution of decoding accuracies for each time point. For each permutation, the time series of decoding accuracies were converted to *z*-values. This was done by comparing each decoding-accuracy value to the respective permutation distribution at the same time point. Continuous clusters with *z*>1.65 (i.e., *p*<0.05) were then identified in each permutation and the respective cluster sums of *z*-values were calculated. From each permutation, the largest cluster sum across all conditions was entered to the maximum-statistic null distribution. Analogously to the conventional procedure [49], each cluster identified from the analysis with true content labels was then compared to this null distribution, to determine their statistical significance. Clusters with *p*_Corrected_<0.05 were considered statistically significant.

### Within-participant Multivariate Pattern Analysis

The within participant analysis is conducted based on the same principle as the cross-participant decoding analysis (Figure 2), by employing the SVM implementation from libsvm [47] and MATLAB/Octave CoSMoMVPA package [48]. The period 50 ms before and after the TMS pulse was not entered into the analysis. Using CoSMoMVPA and Fieldtrip toolboxes in each participant, we first balanced the number of trials for each class and low-pass filtered the signals at 75 Hz. To enhance the SNR, subaverages of four trials were calculated with each class. Twenty different random iterations were calculated of these subaveraged samples. In each iteration, spatial feature selection was performed using a similar, yet individualized, PCA procedure to that used in the cross-participant MVPA (range 5-10 PCs, group median = 8 PCs; see **Suppl. Fig. 1**).

Similar to the cross-participant MVPA, within-participant classification was performed using a temporal searchlight analysis [48] with a 300 ms sliding window in 3 ms steps, conducted separately for the periods before and after the TMS pulse. A *k*-fold cross-validation procedure was used to classify the maintained WM content (*k*=6 in participants with six runs of data; *k*=5 in two participants with five runs of data). In each fold, the model was trained in *k*-1/*k* of the samples and tested in the remaining samples. For each searchlight dataset, the decoding accuracies were averaged across the folds and iterations. For each participant, condition (active TMS, Control TMS), and maintenance period (before or after the TMS pulse), the analyses resulted in a time series with decoding accuracies of each searchlight centroid. Similar to previous studies [30], each participant’s decoding accuracy time courses were smoothed over time with a Gaussian kernel with FWHM of 9.4 ms for significance testing.

### Statistical Significance, Within-participant MVPA

The statistical significance was determined using robust cluster-based permutation testing, which handles multiple comparison problems using a *maximum-statistic* strategy [13]. For each participant and TMS condition, we first generated 500 unique permutations of the true item-content labels of the classifier. The temporal searchlight analysis was repeated with these permuted labels to generate a distribution of decoding accuracies for each time point. To assign a p-value for each time point, the original group-mean decoding accuracy value, found from classifiers with true labels, was compared with this permutation distribution. To improve the precision, we modeled the empirical permutation distribution using a Gaussian fit. Continuous clusters with *p*<0.05 were then identified in each permutation and the respective cluster sums decoding accuracies were calculated. From each permutation, the largest cluster sum across all conditions was entered to the maximum-statistic null distribution. Analogous to the conventional procedure [49], each cluster identified from the analysis with true content labels was then compared to this null distribution, to determine their statistical significance. Clusters with *p*_Corrected_<0.05 were considered statistically significant.

## Results

### Behavioral performance

The participants’ behavioral percent corrects were 79.6 ± 6.5% (mean ± SD) in the Active TMS sessions and 78.7± 6.2% in the Control TMS sessions. Although we see a slight increase in behavioral performance in Active TMS session, the increase is not statistically significant. We additionally calculated the percent correct of answers as a function of the ripple-velocity distance from memory item to probe. This analysis revealed a consistent relationship between participants’ ability to reject non-matching probes and the difference in ripple velocity between the probe and WM item. In the TMS session, the percent correct of answers for match trials was 84.5 ± 6.3%. For non-match trials with JND distance 1, the percent correct was 60.4 ± 13.9%; for JND distance 2, it was 85.7 ± 8.6%, and for non-match trials with JND distance 3, it was 93.9 ± 4.9%. In the Control TMS session, match trial percent correct was 84.1 ± 5.7%. Non-match trial JND distance 1 percent correct was 58.0 ± 14.9%; JND distance 2 percent correct was 2 87.1 ± 10.2%; and JND distance 3 percent correct 92.2 ± 9.0%.

### Multivariate Pattern Analysis Cross-participant MVPA

We conducted a four-class cross-participant classification analysis to determine whether the single TMS pulses to left pSTC enhanced the decoding of memorized content from the ERPs. The analysis employed a temporal searchlight approach, in which the decoding was performed based on the spatiotemporal pattern of EEG activity within a 300-ms sliding window. The statistical significance was verified through a robust cross-participant cross-validation and cluster-based maximum-statistic permutation procedure. According to these analyses, in the Active TMS condition, the MVPA decoding accuracy for memory content rose significantly above chance level during the first few hundreds of milliseconds after the TMS pulse (*p_Corrected_* <0.05, cluster-based maximum-statistic permutation test; cluster sum of normalized accuracy = 137.2; **Fig. 3d**). No statistically significant decoding results were observed in any other time period in the active TMS condition or in the Control condition (for an additional analysis of the stability of decoding, see **Suppl. Fig. 2**).

**Figure 3.**
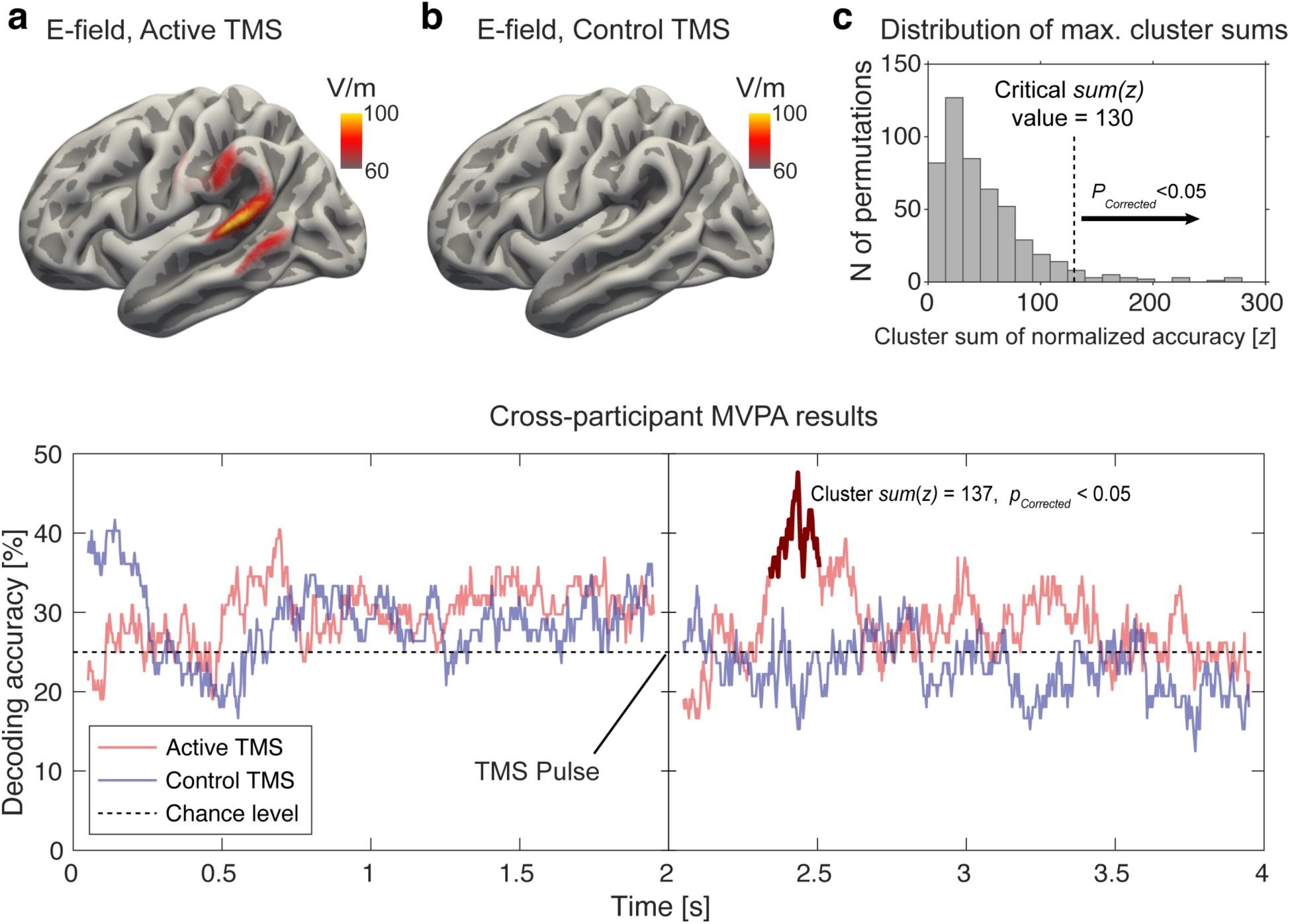
The results of E-field calculations and searchlight MVPA decoding of WM content from EEG. **(a)** Group median E-field maps for the Active TMS condition. **(b)** Group median E-field maps for the Control TMS condition. **(c)** Null distribution for 500 permutations, utilized to determine the statistical significance of decoding accuracies. From each permutation, the maximum cluster sum of normalized decoding accuracy was identified and added to this null distribution. The vertical dotted line illustrates the critical value for *p*_corrected_<0.05 (cluster sum(z) = 130). **(d)** Decoding accuracies in the cross-participant four-class MVPA (% of correctly classified trials). The time series reflect the SVM decoding accuracies at the centroid of each sliding 300-ms searchlight. These decoding accuracies were derived from an iterative leave-one-participant out cross-validation procedure: In each searchlight time window, the data of each participant was used once as the test set and those from the rest of the remaining participants as the training set. The light red line denotes the Active TMS condition and the light blue line the Control TMS condition. The dotted horizontal line indicates the chance level of decoding-accuracy in a four-class classification (25%). The time window when the decoding accuracy was significantly higher than chance level in the Active TMS condition is shown in dark red (*p*_Corrected_<0.05, non-parametric cluster-based permutation test).

### Within-participant MVPA

We additionally conducted a within-participant four-class SVM searchlight to test the effect of the TMS pulse on the individual brain activity. We employed a similar searchlight approach with a 300-ms sliding window as in the cross-participant analysis. Consistent with the cross-participant results, in the Active TMS condition, the group average of MVPA decoding accuracy for memory content rose significantly above chance level during the first few hundreds of milliseconds after the TMS pulse (*p_Corrected_* <0.01, cluster-based maximum-statistic permutation test; cluster sum of decoding accuracy = 24.5; **Fig. 4**). No statistically significant decoding results were observed in any other time period in active TMS or control TMS conditions.

**Figure 4.**
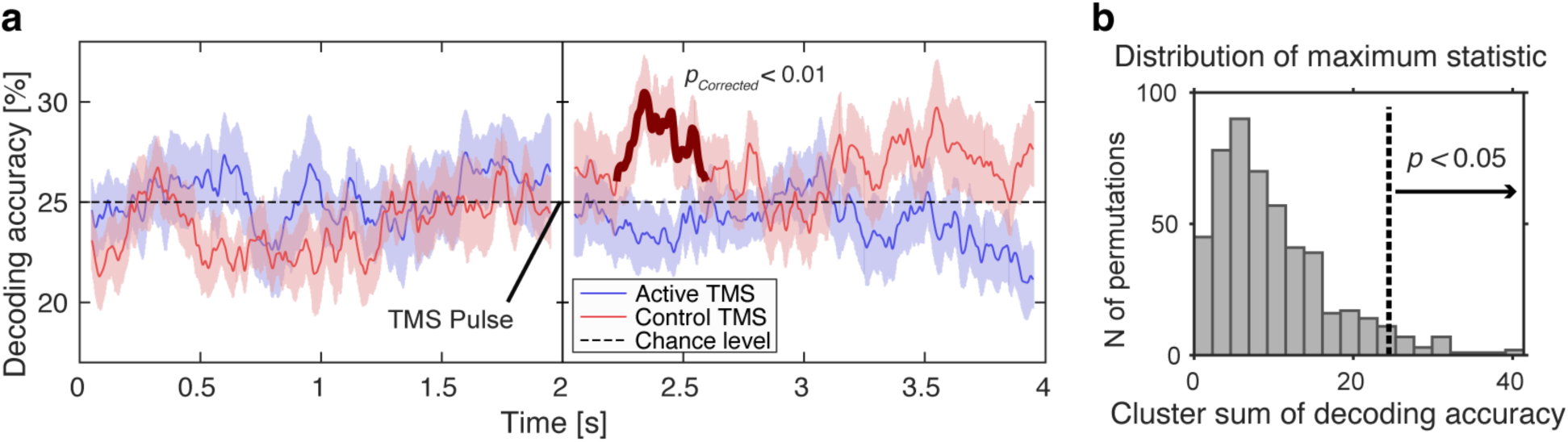
The results of within-participant searchlight MVPA decoding of WM content from EEG for Control and Active TMS conditions. **(a)** Decoding accuracies in the within-participant four-class MVPA. The time-resolved decoding reflects the accuracies at the centroid of each sliding 300-ms searchlight. The thin red line denotes the Active TMS condition and the thin blue line the Control TMS condition. The dotted horizontal line indicates the chance level of decoding-accuracy in a four-class classification (25%). The time window when the decoding accuracy was significantly higher than chance level in the Active TMS condition is shown in dark red (*p*_Corrected_<0.05, non-parametric cluster-based permutation test). **(b)** Null distribution for 500 permutations, utilized to determine the statistical significance of decoding accuracies. From each permutation, the maximum cluster sum of decoding accuracy was identified and added to this null distribution. The vertical dotted line illustrates the critical value for *p*_corrected_<0.05 (cluster sum of decoding accuracy = 24.5).

### E-field Calculations

**Figures 3 a-b** depict the median of the E-field calculations for Active TMS and Control TMS conditions, respectively, thresholded at 60 V/m [50]. The average E-field in the target area (Talairach −60, −38, 15) was 74.11 V/m for the Active TMS condition and 34.75 V/m for the Control TMS condition.

### Control Analyses

To control whether MVPA results are driven by differences in ERP amplitude between different conditions, we averaged the response to different cued content during WM period across participants. No systematic differences were found between signals for different cued content during the maintenance period before or after the TMS pulse. We also performed a grand average of the ERP data to observe the ERP time course during the task trial. **Figure 5** depicts the grand average ERPs calculated for the WM maintenance period as the time of interest and TMS evoked responses (TEP) for Active TMS and Control TMS sessions. As expected, the ERP results revealed a N2/P3 response to the visual retro-cue at the beginning of the maintenance period. A pattern of TMS-elicited ERP deflections was detected 2-2.5 s into the maintenance period. The auditory component reflects a typical TEP elicited by the TMS pulse (**Fig. 5c**). No comparable effect is observed in the Control TMS condition (**Fig. 5d**), although the auditory click sound was identical to the Active condition. This suggests that the click sounds were sufficiently masked by the constant noise masking used in all sessions. Finally, we did not observe any persistent elevation in ERP during the WM maintenance period **(Fig. 5)**. To control the data quality, we calculated grand averages for the item presentation and probe and response period **(Suppl. Fig. 3-4)**. The averaged responses for these periods were as expected.

**Figure 5.**
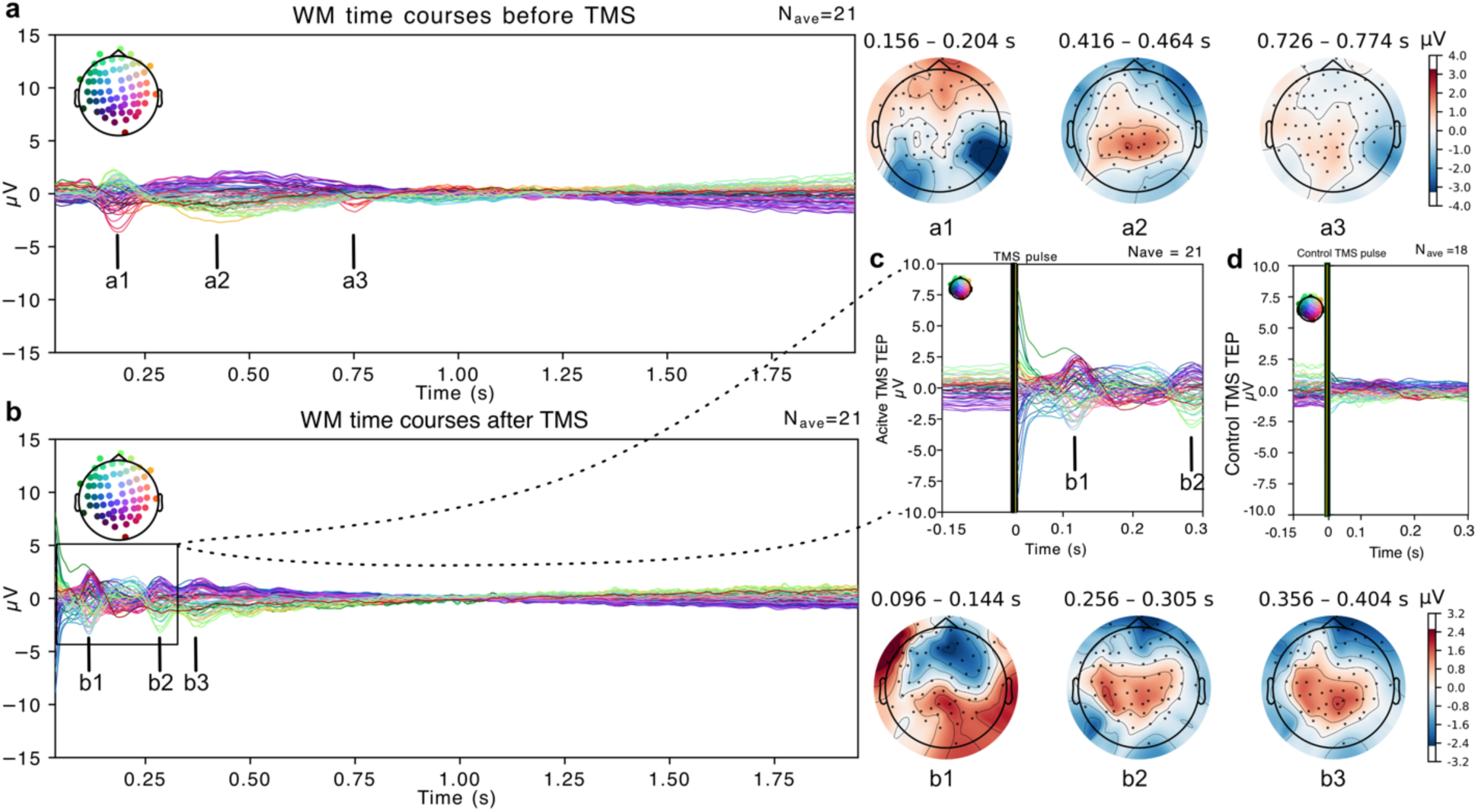
Topographical and butterfly plots of ERP time courses for time of interest (TOI) in the active TMS condition. Different colors in the ERP plots refer to different electrodes. The electrode map above the figures denotes the locations of electrodes. The timeline starts from *t = 0 s* at the visual retro-cue. **(a)** TOI ERP topographical plots and time courses for WM before TMS pulse. The plot depicts the results for the visual retro-cue that starts the maintenance period. **(b)** TOI ERP topographical plots and time courses after TMS pulse. The timeline starts from *t = 0 s* at the TMS pulse. **(c)** TMS evoked response for Active TMS session. **(d)** TMS evoked response for Control TMS session.

To control whether the decoding results were driven by a difference in TEP in different memory conditions due to TMS pulse, we also performed a repeated measures ANOVA across ERPs of different memory conditions. We did not find any significant differences in corrected or uncorrected level between averaged ERPs across different conditions (35-100ms: F_3,60_=0.65, p=0.59; 100-200ms: F_3,60_=0.22, p=0.88; 200-300ms: F_3,60_=0.36, p=0.78). We also tested the ERPs across the memory period after the TMS pulse to test whether the auditory click artifact might bias our main analysis results. We found no significant differences in the trial-averaged EEG patterns following the TMS pulses across the memory conditions during the whole 2-s memory period after the TMS pulse (F_3,60_=0.28, p=0.84).

Finally, we also performed a control decoding analysis with the same parameters as our main analysis using the task-irrelevant (i.e., “un-cued”) items, which were to be forgotten after the retro-cue. This control MVPA showed no significant decoding for neither Active TMS nor Control TMS conditions at any time during the WM retention period.

## Discussion

Here, we investigated auditory WM using a “perturbation approach”, which combines MRI-navigated single pulse TMS with simultaneous EEG recordings, to unravel content-specific mnemonic states from EEG otherwise obscured by the much larger EEG “background” activity. To decode WM content from EEG signals during the maintenance period, we employed a temporal-searchlight MVPA with robust cross-participant cross-validation and non-parametric permutation testing. As predicted, the decoding accuracy of auditory WM content rose significantly above chance level after a single TMS pulse was delivered to the non-primary auditory areas in the left pSTC. Further, the Control TMS condition (otherwise the same as active single pulse TMS but with a 20-mm plastic block between the coil and the participant’s scalp) found no significant decoding accuracy.

One possible explanation for the present finding is offered by the synaptic theory of WM [26]. According to this model, WM information is coded to content-specific patterns of functional connectivity, which result from activity-based STSP in the synaptic terminals of neurons that are strongly activated at the encoding stage [26]. Instead of persistent neuronal activity, this model predicts that only sparse bursts of neuronal oscillations and firing activity are necessary to maintain this otherwise activity-silent mnemonic brain state [3, 51–53]. Notably, although synaptic states are not directly measurable by non-invasive recordings, computational modeling predicts that a circuit that maintains information by content-specific changes of synaptic weights responds differently to other impulse stimuli (until the STSP decays) [26]. These content-specific responses to external impulses might not only provide a way for the maintained content to be read out at the recall stage, but they could also allow one to “ping” the maintained content with externally generated pulses such as single pulse TMS [13, 28–30].

Whereas the original synaptic theory refers to local circuits in prefrontal cortices [26], it has also been proposed that this synaptic model can also provide a way to explain how WM information might be represented in sensory areas, which presumably cannot support persistent neuronal firing in the absence of sensory stimulation [54–56]. However, previous studies probing activity-silent states of WM have been limited to TMS-based perturbation of association areas [28], or to using auditory or visual “impulse stimuli” [13, 31] that might activate a wide array of brain networks beyond sensory areas involved in orienting to task-irrelevant stimulus changes. TMS offers a more direct way to focally perturb cortical neurons [33, 57] therefore increasing the likelihood that the content-related signals originate from the targeted sensory areas rather than from higher-order brain regions. The present results suggest that WM content can be decoded at a high accuracy from EEG responses to TMS pulses directed to pSTC. They could thus offer new insights into the role of activity-silent WM processes in the sensory cortices.

Alternative explanations for enhanced decodability of WM content, which follows an irrelevant impulse stimulus (i.e., “pinging effects”), have been recently presented. In a recent reanalysis of previous studies [28, 30], Barbosa et al. [58] attributed “pinging effects” of visual WM content to reduced trial-to-trial variability of EEG signals, which was observed after the strong visual impulse stimuli that had been used to facilitate the decoding of (presumably activity-silent) WM content (see also [59]). Their reasoning was that such a reduction of variability across trials could have enhanced the performance of algorithm because of enhanced SNR, rather than a genuine WM reactivation effect. It is, however, important to note that there were notable differences between the pinging effects of visual impulses vs. TMS-induced perturbations. In contrast to the effect of strong visual impulse stimuli (reanalysis the data in [60]), TMS perturbations increased, rather than decreased, the variability of signals from trial-to-trial (reanalysis of [28]). *These results led Barbosa et al. to conclude that TMS-induced enhancement of WM decoding from EEG data could, nonetheless, reflect an activity-silent mechanism of WM. Notably, consistent with Barbosa et al., the present analyses provide no evidence of TMS-induced reduction of trial-to-trial variability of EEG signals, which could have explained the enhancement of the decoding accuracy of auditory WM content*.

Using measures such as EEG to probe synaptic processes is supported by the notion that EEG signals primarily result from post-synaptic processes in apical dendrites of cortical pyramidal neurons [61–63]. These neurons constitute fundamental components of the canonical cortical circuit that presumably supports WM [25, 64–66]. An inherent limitation of non-invasive measures, however, is that they do not conclusively rule out other alternative explanations. In addition to an activity-silent synaptic state, the present enhancement of WM decoding by TMS pulses could also result from perturbation of an activity-based maintenance process. Cellular-level studies demonstrate that single pulse TMS activates a broad population of cell bodies in the cortex [57, 67]. The rapid firing of these neurons after the TMS pulse, which is followed by a refractory period, disrupts the cortical network, resetting the stimulated region [57, 67]. The present results could therefore also be arguably consistent with a subthreshold attractor model [59, 68]. The subthreshold attractor model suggests that WM-related persistent activity tends to be attracted to a bump state that emerges in varying locations across this network [69, 70]. By interfering with such activation patterns, TMS pulses might result in content-specific signal changes that are recordable by EEG. However, a challenge for such a model in the present context is that the TMS pulse, which tends to overwrite the neuronal activity in the stimulated area, would disrupt the content-specific population activity in the stimulated network. Neurophysiological studies provide experimental evidence indicating that distractor events disrupt content-specific firing activities rather than amplifying them to a discernible level in the mass action of neurons [71]. Therefore, while it is not entirely incompatible, the subthreshold attractor model does not adequately describe our results because we found no evidence of impaired WM performance in the Active TMS vs. Control TMS condition.

Another alternative for a hidden state (whether it is due to synaptic plasticity or to other means) of WM content in sensory areas is that the WM maintenance is carried through a recurrent neural network where the PFC shapes and transforms the WM representations according to task demands [54, 72]. The recurrent model is another possible explanation for how the synaptic weights could be formed and maintained in the posterior and sensory brain areas. Indeed, it has been recently shown that the WM content can be effectively maintained by a neuronal behavior explained by a combination of activity-based and activity-silent models of WM [73].

Another important consideration is that, although the initial E-field exceeded the stimulation threshold only in our targeted STG site, single-pulse TMS could influence not only local but also distant neural circuits through axonal and transsynaptic propagation, potentially affecting other cortical and subcortical areas starting already at the first tens of milliseconds after the TMS pulse.

The action potentials generated by the TMS-induced electric field may propagate along the axons in both anterograde and retrograde directions, facilitating forward and backward information flow within the stimulated pathway [74]. Computational modeling studies of TMS-EEG effects suggest that recurrent network feedback to the target regions begins driving TEP responses already around 100 ms post-stimulation, whereas only the earlier TEP components can be attributed to local reverberatory activity within the stimulated region [75]. Roughly consistent with previous impulse-stimulus and TMS studies of visual WM [28, 30], here the significant increases of decoding accuracy occurred about 200-300 ms after the TMS pulse, peaking slightly earlier in the within-participant than cross-participant analyses. The observed effects may result from feedforward propagation of activity from STG to other areas and/or subsequent feedback influences from other regions back to STG.

Our cross-participant decoding results indicate that the states that were revealed by the TMS pulse were stable across participants[76]. This is argued to be similar to the difference between fixed- and mixed-effects analyses [77]. In the context of MVPA, this distinction allows for the identification of differences in local computations. Significant prediction in a cross-participant model indicates that the stimulus-related information encoded by a time-resolved neuronal activity pattern stays relatively consistent across participants [78].

Some inherent limitations of EEG interpretations during a combined TMS-EEG study include several types of artifacts such as direct muscle/sensory nerve stimulation, somatosensory sensation related to the vibration of the coil, and acoustical clicks. There is a possibility that the improved decoding following a TMS pulse is attributable not only to its neurophysiological effects on the target area but also to non-specific effects associated with unrelated physiological events. Here, we attempted to control these biases with a Control TMS condition. Adding a hard plastic block provides similar tactile and auditory sensation as Active TMS but with subthreshold brain stimulation. To mitigate the effect of acoustical clicks we also used a continuous noise masker stimulus and TMS compatible insert earphones accompanied with earplugs that attenuate the background noise [37]. It is also worth noting that the same TMS stimulator output level was used for Control and Active TMS sessions for each participant, resulting also in exactly the same sound level of the TMS click sound. Our continuous noise masking should have mainly eliminated the possibility of TMS-evoked auditory effects (**Fig. 5 a-b**). Further, such effects should be similar across the control and active TMS conditions as well as between the different memory conditions. It is thus unlikely that the decoding results would be biased by any auditory artifacts. The differences in WM decoding between Active and Control sessions thus cannot follow from the click sound, per se. Finally, TMS was applied at a fixed 2-s latency, which might have created an anticipation effect However, such an anticipation effect should have been identical in the Active TMS and Control TMS conditions, making it unlikely that our main results were influenced by such an effect.

To conclude, using TMS-EEG and cross-participant MVPA, the present study suggests maintenance of WM content involves an “activity-silent” brain state in auditory brain areas. The study also demonstrates the power of TMS as a way to probe information content embedded in EEG signals.

## Acknowledgments

This project was supported by NIH grants R01DC016915, R01DC016765, R01DC017991, 1R01DC020891, R01MH128421, R01MH130490-01A1, R01NS126337-03, R01NS126337, S10OD028668 and P41EB030006.

## Open practices statement

The deidentified data and code to reproduce the main findings will be made available on https://dataverse.harvard.edu/dataverse/isiluluc/.

## Declaration of Interest

Aapo Nummenmaa and Lucia Navarro de Lara are named inventors in patents and patent applications related to TMS.

Tommi Raij and Mohammad Daneshzand are named inventors in patent applications related to TMS.

## Notes

### Competing Interest Statement

The authors have declared no competing interest.

### Summary of Updates

This revision includes additional analysis and results to the existing manuscript.

