## Supplementary material for "Decoding auditory working memory content from EEG aftereffects of auditory-cortical TMS"

### Supplemental Material

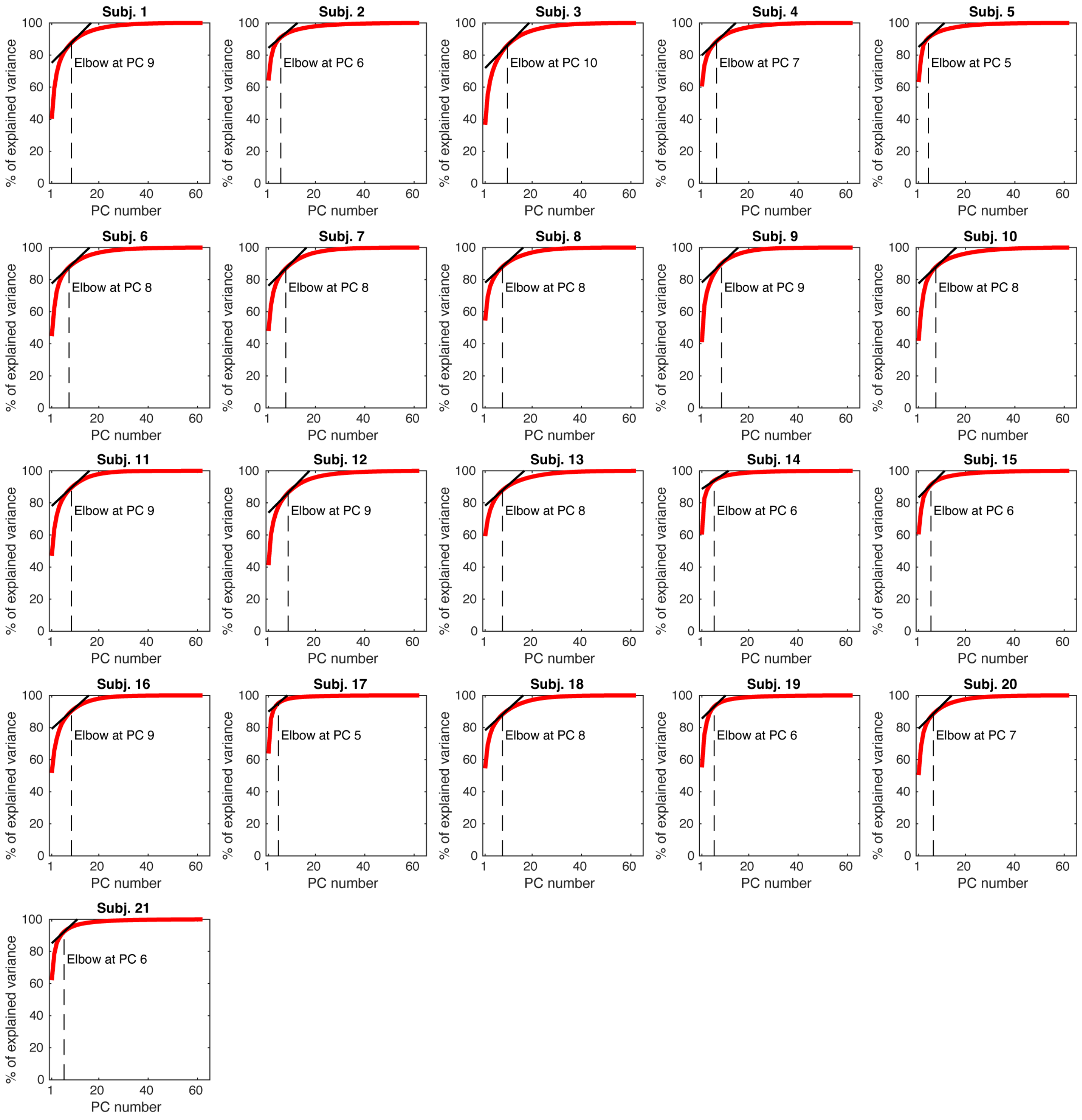


**Supplementary Figure 1.** Selection of the optimal number of principal components (PCs) using the "elbow" method in each individual participant. The cumulative variance explained by each successive principal component is plotted, with the "elbow" point indicating the optimal number of PCs.

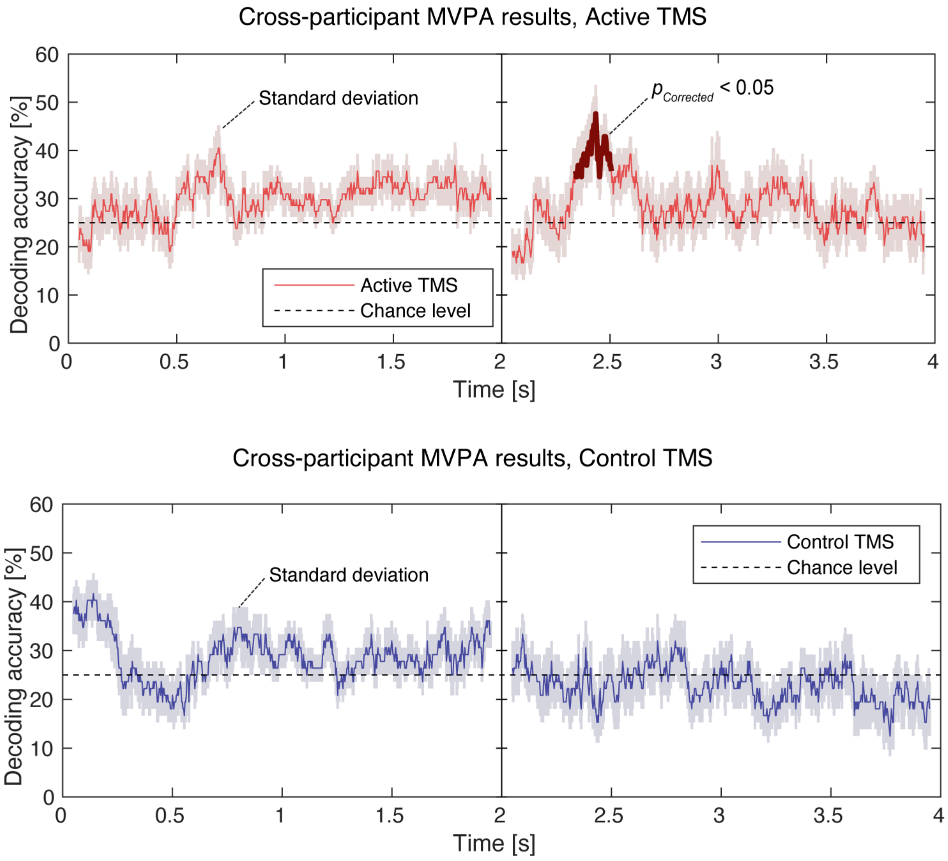


**Supplementary Figure 2.** Results with bootstrapping to obtain the standard deviation (SD) of the cross-participant MVPA. The shaded areas depict the bootstrapping SD for Active TMS (upper) and Control TMS (lower) sessions. The time window when the decoding accuracy was significantly higher than chance level in the Active TMS condition is shown in dark red ($p_{Corrected}$<0.05, non-parametric cluster-based permutation test).

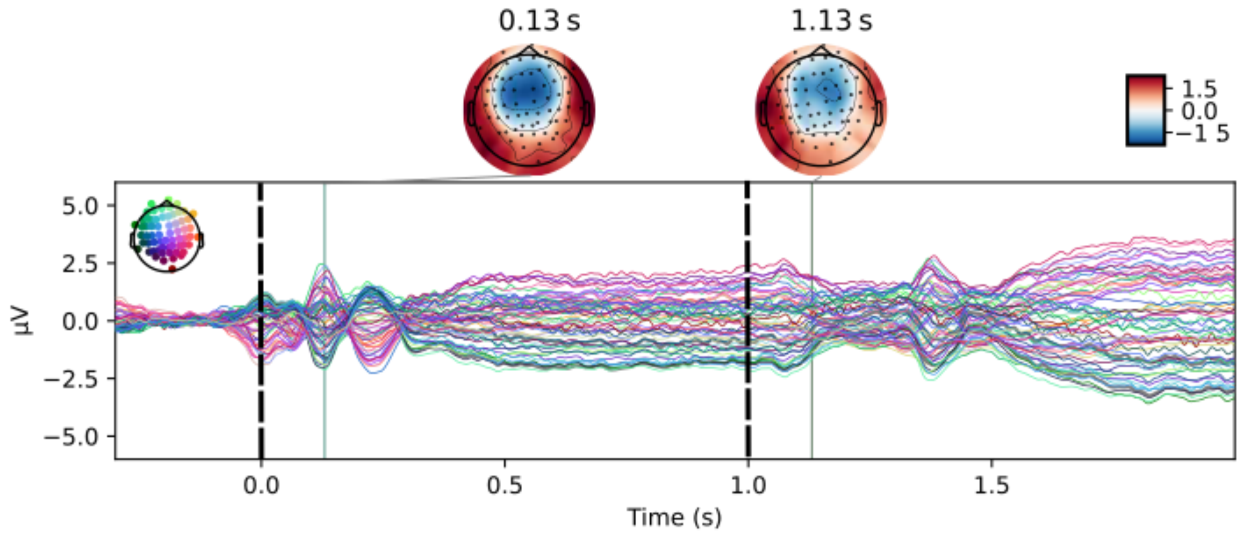


**Supplementary Figure 3.** ERP responses to the presentation of ripple-sound working memory (WM) items. The ripple sound onsets are denoted with black dashed lines. The onset of the first ripple sound is *t=0 s*. Onset of the second ripple sound item is *t=1 s*. The different colors on the butterfly plot and the map on it denote different electrodes. The left topographical map (t=0.13 s) demonstrates the scalp potential distribution at the peak of the N1 response to presentation of the first auditory WM item. The right topographical map (t=1.13 s) shows the scalp potential distribution at the corresponding latency after presentation of the second auditory WM item.

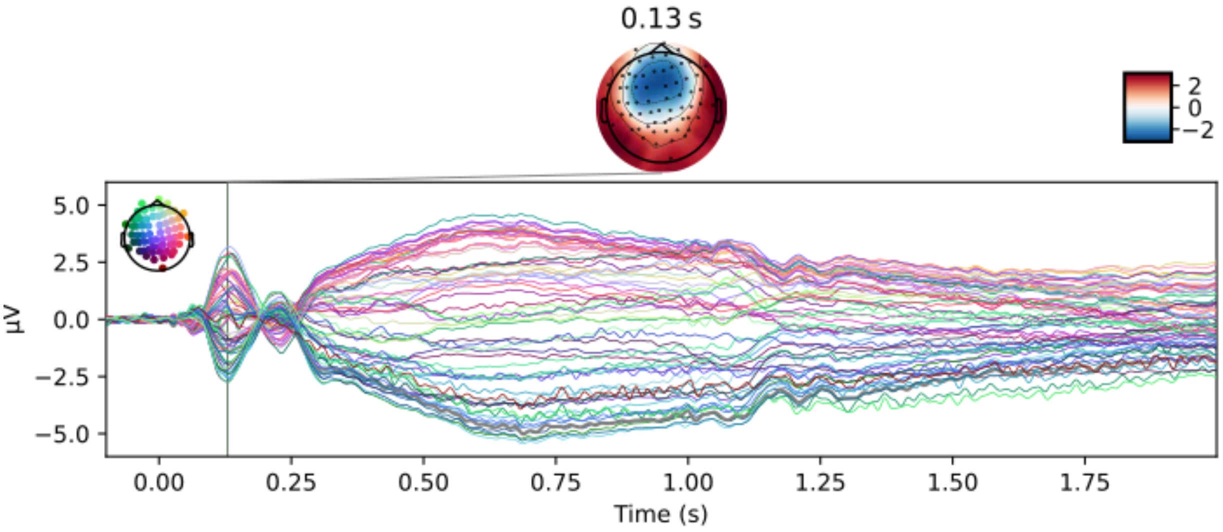


**Supplementary Figure 4.** ERP responses to the presentation of WM probe and during the recall period. The onset of the probe ripple sound is t=0 s. The scalp potential distribution is shown at the peak of the N1 response to test sound item.
